# Modulating short-term auditory memory with focal transcranial direct current stimulation applied to the Supramarginal Gyrus

**DOI:** 10.1101/2021.01.27.428474

**Authors:** Karl D. Lerud, Bradley W. Vines, Anant B. Shinde, Gottfried Schlaug

## Abstract

Previous studies have shown that transcranial direct current stimulation (tDCS) can affect performance by decreasing regional excitability in a brain region that contributes to the task of interest. To our knowledge, no research to date has found both enhancing and diminishing effects on performance, depending upon which polarity of current is applied. The supramarginal gyrus (SMG) is an ideal brain region for testing tDCS effects because it is easy to identify using the 10-20 EEG coordinate system, and results of neuroimaging studies have implicated the left SMG in short-term memory for phonological and non-phonological sounds. In the present study, we found that applying tDCS to the left SMG affected pitch memory in a manner that depended upon the polarity of stimulation: cathodal tDCS had a negative impact on performance while anodal tDCS had a positive impact. These effects were significantly different from sham stimulation, which did not influence performance; they were also specific to the left hemisphere – no effect was found when applying cathodal stimulation to the right SMG – and were unique to pitch memory as opposed to memory for visual shapes. Our results provide further evidence that the left SMG is a nodal point for short-term auditory storage and demonstrate the potential of tDCS to influence cognitive performance, and to causally examine hypotheses derived from neuroimaging studies.

## 1 Introduction

Neuroimaging data support a critical role for the left supramarginal gyrus (SMG) in short-term storage of verbal (1–4) and non-verbal auditory information (5–10). In previous studies, our group has found that the degree of activation in the left SMG is correlated with performance on a pitch memory task (6), that SMG activation also changes over time in a pre-post design of a short-term auditory memory learning study, and that the amount of activation change correlates with the improvement in the auditory learning experiment (7). Based upon these neuroimaging studies, we may infer that pitch memory performance is correlated with activity in the left SMG. Employing non-invasive brain stimulation allows us to test hypotheses about brain-behavior relations, and to directly assess causality.

Transcranial Direct Current Stimulation (tDCS) is a non-invasive brain stimulation method that uses a weak electrical current to modulate the excitability of brain tissue. The effects of tDCS vary with current strength, stimulus duration, and the direction of current flow, as determined by electrode size, position, and polarity (anodal vs. cathodal) (11–18). Several studies have shown that applying tDCS can impact performance on tasks that involve the brain region receiving stimulation. Researchers have found that applying cathodal tDCS can impair performance on certain tasks (8,19–25), whereas anodal tDCS can enhance performance (21,26–30). For example, our group (21,31) found evidence that applying anodal tDCS over the left pre-central gyrus (C3 of the international 10-20 system) could improve the accuracy of contralateral (right) finger-sequence movements, while other groups have also found evidence for improvements in the auditory domains (9,30,32). However, to our knowledge, no tDCS study has found significant effects on cognitive performance depending on the polarity of stimulation, in which there would be an improvement in performance with anodal tDCS and a deterioration in performance with cathodal tDCS. In comparison to (9,30,32), we are testing whether or not subjects can remember the pitch of an initial note by comparing it to the last note in a sequence that contains distractor tones, similarly to classic pitch memory experiments (33–35), while other (9,30,32) studies utilized a same/different comparison of two melodies. Furthermore, we are using the exact experiment that revealed a specific engagement of the supramarginal gyrus in a previous functional imaging study on pitch memory (6,7) .

In this paper, we present two experiments investigating the causal influence of the left SMG on short-term memory for pitch. The first experiment replicated our previous study (8), in which we investigated how applying cathodal tDCS over the left SMG affected pitch memory performance, compared with sham tDCS and cathodal tDCS over the right SMG. In the second experiment, we compared anodal and sham tDCS, as applied over the left SMG, and measured performance on both a pitch memory task and a visual memory task that we designed to control for the possibility of a domain-general effect. Our hypothesis was that applying cathodal tDCS over the left SMG would have a negative impact on pitch memory performance, whereas applying anodal tDCS would have a positive impact. We also anticipated that the tDCS would only affect pitch memory (as opposed to visual memory), and that only stimulating the left SMG would have a significant effect on performance (as opposed to stimulating the right SMG). Such outcomes would not only underscore the importance of the left SMG to short-term storage of pitch information, but would also highlight the potential of transcranial direct current stimulation as a tool for investigating causal relations between brain areas and cognitive operations.

## 2 Materials and methods

### 2.1 Experiment 1: Cathodal tDCS – controlling for location of stimulation

#### 2.1.1 Participants

Fourteen (14) healthy participants (7F, 7M) took part in experiment 1, after giving their informed written consent in accordance with the Beth Israel Deaconess Medical Center ethics review board. Musical ability or training was not used as a screening criterion. Most of the participants had some musical training, but none were actively pursuing instrumental music training at the time of the experiment, and none had been a professional musician. All subjects reported having normal hearing and it was required that subjects heard the test tones at equal loudness in each headphone, otherwise subjects would have been excluded.

#### 2.1.2 Pitch memory task

This task was identical to that used in the (6,7) neuroimaging studies, and in our previous study (8). There were 39 stimuli. Each stimulus comprised a sequence of sine-wave tones. The tones were all 300 ms in duration, of the same amplitude, and separated by a constant 300 ms inter-stimulus interval. The frequencies used for targets were taken from the semitones of the Western chromatic scale, in the range from E4 (330 Hz) to D#5 (622 Hz). The distractor tones were microtones within the same frequency range. There were six tones in one half of the sequences (one target, four distractors, one target), and seven tones in the other half (one target, five distractors, one target). The task instructions for a single trial were to register as quickly as possible whether the first and last tones in a sequence were the same or different. One mouse button corresponded to the answer “same” and the other to “different”. The 39 pitch sequences were presented in a new randomized order for each iteration of the task. The task lasted approximately 5.2 minutes. Participants wore noise-cancellation headphones, and were instructed not to sing or hum during the task performance.

#### 2.1.3 Procedure

The experimental procedure involved one day of practice and two days of testing. The practice session took place on the day prior to the first day of testing. Participants practiced the pitch memory task until reaching a plateau in performance; we set the threshold for this stable performance level as 1.) having completed at least three practice runs, and 2.) having changed less than two points in performance over the last two runs. On the first day of testing, participants performed one warm-up run of the pitch memory task, and then underwent two tDCS sessions, one for cathodal stimulation over the left SMG, and one for sham tDCS; the ordering of cathodal and sham stimulation was counterbalanced across participants. For each tDCS session, participants completed one pre-stimulation run at baseline, and one post-stimulation run immediately following 20 minutes of stimulation. The two tDCS sessions (cathodal and sham) were separated by a washout period of at least 60 minutes to avoid carryover effects of the stimulation. We found this to be a sufficient duration in our own previous experiments (8,21,31) and according to the literature; though studies have found that physiological effects from tDCS can last for 90 minutes beyond the period of stimulation (36,37), to our knowledge there have been no reports of effects on behavior lasting longer than 30 minutes after a single session of tDCS (20,38,39). The anodal elliptically-shaped electrode (area = 16.3 cm^2^) was centered over CP3 of the international 10–20 system for electroencephalography (EEG) electrode placement, which corresponded to the left SMG and contains Brodmann area 40 (40). We conducted a pilot study (n=5) with high resolution MRI (1 mm^3^ voxel size) to confirm that CP3 corresponds to the left SMG; Fig. 1 shows a surface reconstruction from a T1 scan for a representative subject. The cathodal electrode (rectangular, area = 26 cm^2^) was secured over the contralateral supraorbital area, and was functionally ineffective in this experimental design (41,42) .

**Fig 1:**
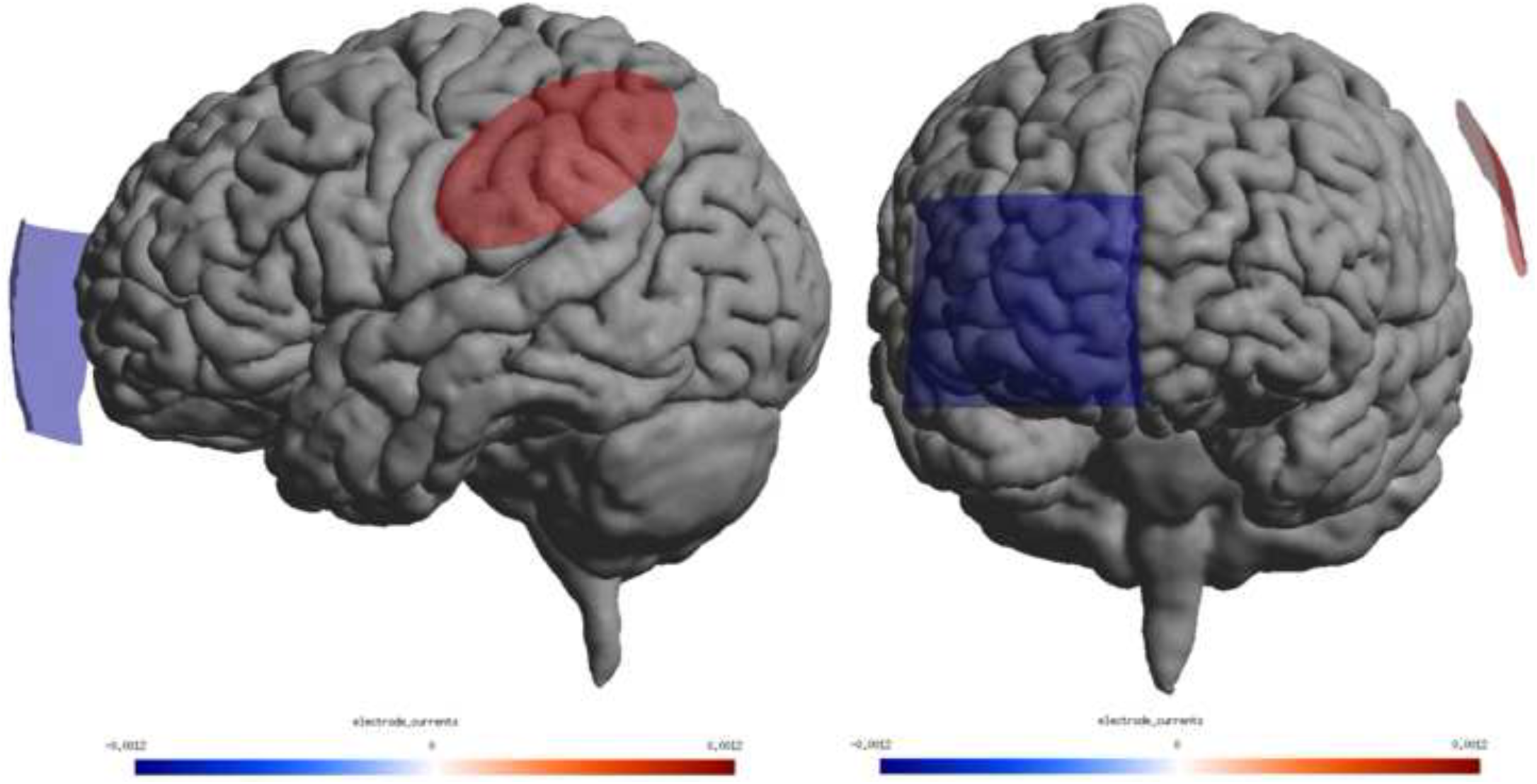
Surface reconstruction from high-resolution T1 scan obtained with SimNIBS, along with representative electrode locations superimposed. In this case, the SMG electrode was the anode, shown in red.

The second day of testing took place at least one week after the first day of testing. We included the second day of testing to determine whether there was a hemispheric asymmetry in the effects of tDCS on pitch memory. Participants underwent cathodal stimulation over the right SMG. The cathodal electrode was centered on CP4 and was the same elliptical electrode that was previously the anode, corresponding to Brodmann’s area 40 in the right hemisphere. The anodal electrode was positioned over the left supraorbital area and was the same rectangular shape as when it served as the cathode previously.

For the real stimulation sessions (cathodal stimulation over the left or right SMG), a constant current stimulator (Phoresor II PM850; Iomed Inc., Salt Lake City, Utah, USA) delivered a 1.2 mA current for 20 minutes. For sham stimulation, the current was allowed to ramp up over the first 30 seconds before the experimenter reduced the current to 0, and it remained at 0 for the remaining time period. Participants reported a tingly/itchy sensation at the start of the stimulation, which typically faded away after a few seconds. This sensation was the same for real and sham stimulation. (43) found that naive participants were not able to distinguish between real and sham tDCS, as employed in a manner similar to the present study.

#### 2.1.4 Data analyses

We entered the data into a 3 by 2 repeated-measures ANOVA, with factors tDCS condition (sham, cathodal over left SMG, cathodal over right SMG) and time (pre-stimulation, post-stimulation) with the dependent variable the number of correct answers in the pitch memory task. Post-hoc analyses with a Bonferroni correction compared baseline performance across conditions, as well as effects from pre-to post-stimulation within conditions.

We also calculated the proportion of change in the number of correct trials from pre-stimulation to post-stimulation for each condition, and ran a one-way repeated-measures ANOVA with three levels (sham, cathodal over left SMG, cathodal over right SMG) on the outcomes. Post-hoc analyses with a Bonferroni correction compared the results for the three conditions directly. We used the proportion of change, instead of a simple difference (post - pre), in order to control for variation in skill level across the participants.

### 2.2 Experiment 2: Anodal tDCS - controlling for perceptual domain

#### 2.2.1 Participants

Twelve (12) healthy right-handed volunteers (age range: 21-42 years; 6 male) gave their informed, written consent to participate in this study, which was approved by the local ethics board. None of the subjects from Experiment 1 took part in Experiment 2. The average number of years of formal musical training for the participants was 5.8 years (SD = 5.9); none of the participants were actively practicing a musical instrument at the time of testing; none were music students at a conservatory or professional musicians. All participants reported having normal hearing and normal visual acuity. They all scored within the normal range of intelligence, as measured with the standardized Raven’s Advanced Progressive Matrices test (APM) (44) .

#### 2.2.2 Tasks

We used both the pitch memory task and a visual memory control task in this study. The pitch memory task was identical to the one used in Experiment 1. The visual memory control task was similar in design to the pitch memory task (see Fig. 2). The stimuli were 39 series of pictures. Each picture was an assembly of small black squares (figure height = 5 cm, width = 4 cm), presented in a 6 cm by 6 cm presentation square, on a 30 cm by 32.5 cm computer monitor (15 inch). The monitor was located approximately 60 cm from the seated participants. Each picture was presented for 300 ms, with a blank screen appearing for 300 ms between consecutive pictures. In the context of a sequential array of images, previous research suggests that it takes 200-300 ms of focused attention to consolidate one display in visual short-term memory as a stable representation (45,46). All of the participants reported that they perceived each picture as a separate image, embedded in a series of pictures. The instructions for a single trial were to register as quickly as possible whether the first and last pictures in a picture series were the same or different, by pressing one of two mouse buttons, as in the pitch memory task. The 39 picture sequences were presented in a new randomized order for every iteration of the task.

**Fig 2:**
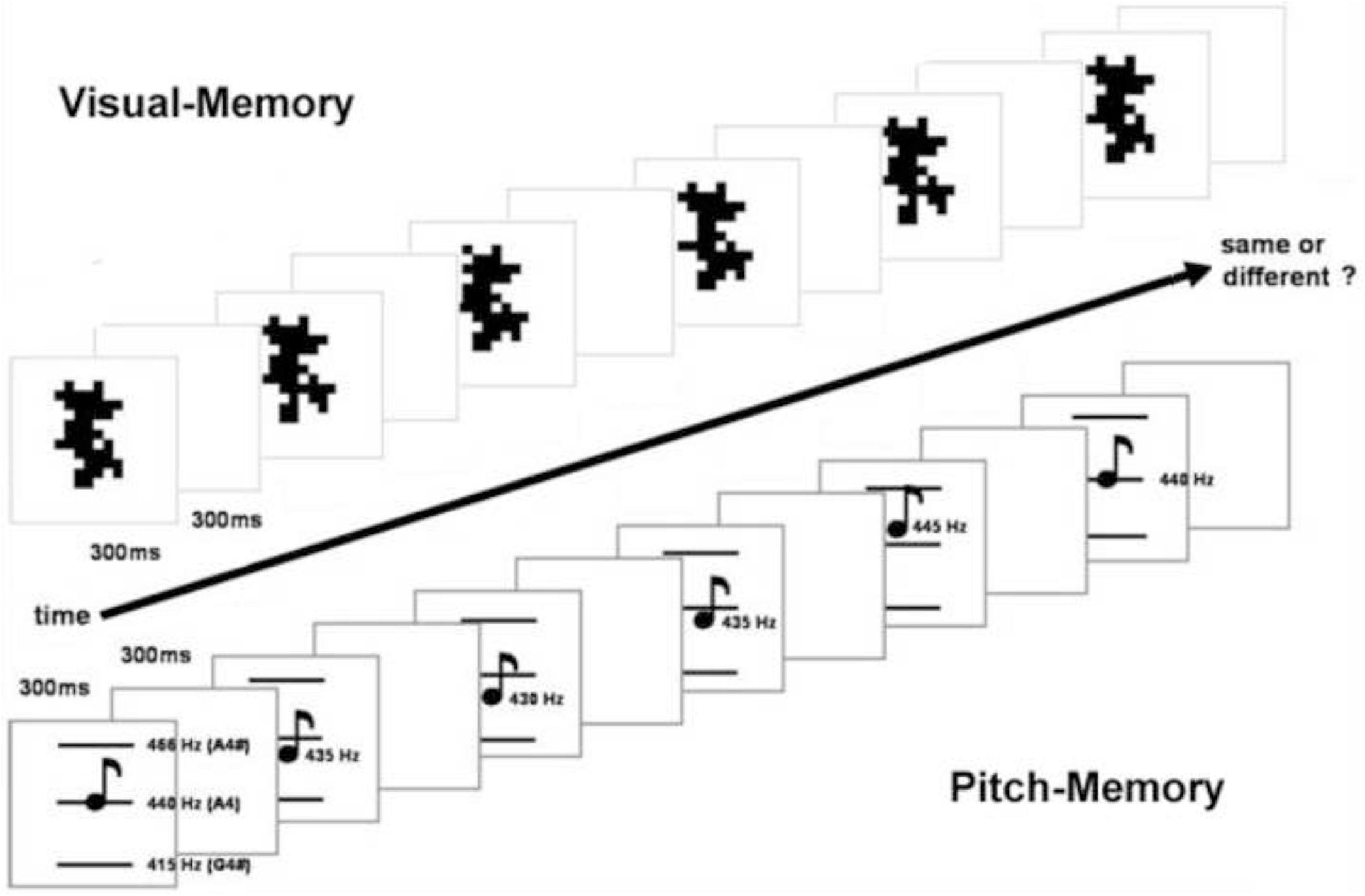
Illustration of experimental procedure for both the pitch memory and visual memory tasks.

We ran a pilot study with 5 participants to determine if the level of difficulty differed between the pitch and visual memory tasks and we used the pilot subjects experience to optimize the experimental setup to ensure that both tasks could be performed with equal accuracy. These pilot experiments differed both qualitatively and quantitatively from the real experiments reported in this paper and are therefore not included in the analysis.

#### 2.2.3 Procedure

The experiment involved one day of practice and two days of testing. During the day of practice, we assessed participants’ handedness, and collected questionnaire data on musical training and other background information; the participants also became familiar with the tasks by performing each task two times. We found in Experiment 1 that two practice runs was sufficient to achieve a stable plateau in performance.

On each experimental testing day, participants completed a warm-up run consisting of just 10 stimulus sequences for the pitch and visual memory tasks. During the warm-up run, participants adjusted the volume and their distance from the screen to comfortable levels, which were then held constant through the course of the session. Each experimental session included a pre-stimulation test, followed by 20 minutes of anodal or sham tDCS stimulation, and a post-stimulation test immediately following the application of tDCS. Pre- and post-stimulation tests included one run of the pitch memory task and one run of the visual memory task. The ordering of the two tasks was counterbalanced across participants and held constant for the two days of testing within participants. On one testing day, sham tDCS was applied, whereas anodal tDCS was applied on the other testing day. The ordering of sham and anodal stimulation was counterbalanced across participants.

#### 2.2.4 Transcranial direct current stimulation

Saline-soaked electrodes were placed on the scalp as in the left SMG condition from Experiment 1. The electrode which was either anodal or sham (area = 16.3 cm^2^) was centered on the left CP3, while the cathodal electrode (area = 30 cm^2^) was positioned over the contralateral supraorbital area. A battery driven constant-current stimulator (Phoresor ® II PM 850; Iomed Inc., Salt Lake City, Utah) delivered 1.2 mA of current for 20 minutes in the anodal tDCS condition. The sham stimulation was administered as in Experiment 1. When asked explicitly, participants did not report having felt any differences between the real and sham sessions.

#### 2.2.5 Data analysis

The pitch memory and visual memory data were entered into separate statistical analyses. We applied 2 by 2 repeated-measures ANOVAs, with factors time (pre-stimulation, post-stimulation) and condition (sham, anodal). Post-hoc analyses with a Bonferroni correction compared baseline performance across conditions, as well as effects from pre-to post-stimulation within conditions. We also calculated the proportion of change in the number of correct answers from pre-stimulation to post-stimulation for each condition, and ran a paired-samples t-test comparing the two conditions (sham, anodal tDCS over the left SMG).

## 3 Results

### 3.1 Experiment 1: Cathodal tDCS – controlling for location of stimulation

We applied a 3 by 2 repeated-measures ANOVA to the data, summarized in Fig. 3. The dependent variable was the total number of correct trials during the pitch memory task, and the ANOVA factors were tDCS condition (cathodal over the left SMG, cathodal over the right SMG, and sham), and time (pre-stimulation, post-stimulation). There were no main effects, but there was a significant interaction between the factors tDCS condition and time (*F(2,26) = 6*.*17, p = 0*.*006*). This provided evidence that the relation between performance pre- and post-stimulation depended upon the tDCS condition. We ran Bonferroni-corrected post-hoc comparisons to determine if performance before stimulation differed from performance after stimulation for any of the tDCS conditions individually, and to compare baseline (pre) and post-stimulation performance across the three tDCS conditions. The comparisons of pre- and post-stimulation performance for each stimulation condition revealed that only cathodal stimulation over the left SMG had a significant effect (*F(1,13) = 25*.*4, p = 0*.*0007*); there was a significant decrease in performance due to applying cathodal tDCS over the left SMG. Pre- and post-stimulation performance did not differ for sham (*F(1,13) = 0*.*515, p > 0*.*9*) or for cathodal stimulation over the right SMG (*F(1,13) = 1, p > 0*.*9*). The post-hoc analysis to test whether or not there was an effect of condition on either the pre- or post-stimulation time periods revealed a trend for an effect for the baseline, pre-stimulation period (*F(2,26) = 4*.*03, p = 0*.*06*). This effect was not significant, thus pairwise tests were not run. There was not a difference between post-stimulation performance for the three conditions (*F(2,26) = 2*.*72, p = 0*.*168*). The baseline comparisons established sham as a valid control condition for cathodal tDCS over both the left and the right SMG.

**Fig 3:**
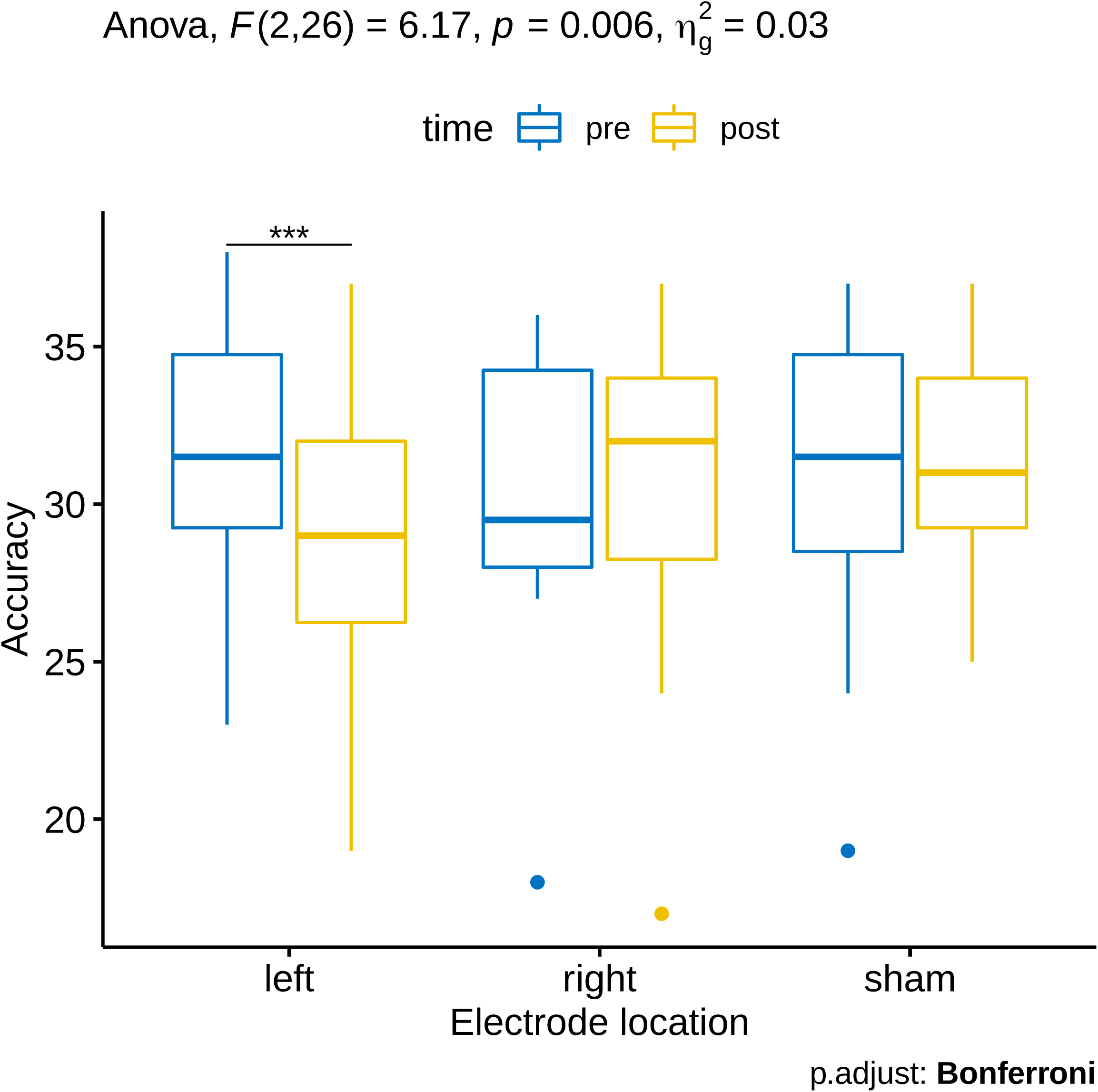
Repeated measures ANOVA and post-hoc comparisons revealed an interaction between the factors electrode condition (left cathodal, right cathodal, sham) and time (pre and post stimulation). A corrected post-hoc test showed a significant decrease in performance for left cathodal stimulation, but not for right or sham stimulation.

We also analyzed the data with a one-way repeated measures ANOVA with three levels for the three tDCS stimulation conditions. The dependent variable was the proportion of change in performance from before to after tDCS. This analysis yielded a main effect of tDCS condition (*F(2,26) = 5*.*63, p < 0*.*01*), which established that there was a significant difference between the effects of at least two of the conditions. Fig. 4 shows the data and associated ANOVA results for proportion of change. A post-hoc analysis with Bonferroni correction revealed significant differences between the effects of cathodal tDCS over the left SMG and both of the other conditions: sham (*p = 0*.*046*), and cathodal tDCS over the right SMG (*p = 0*.*002*). The effects of sham and right-hemisphere stimulation did not differ (*p > 0*.*9*). This analysis revealed that only stimulating the left SMG with cathodal stimulation led to a significant decrease in performance compared to sham. There was no such effect associated with stimulating the right SMG.

**Fig 4:**
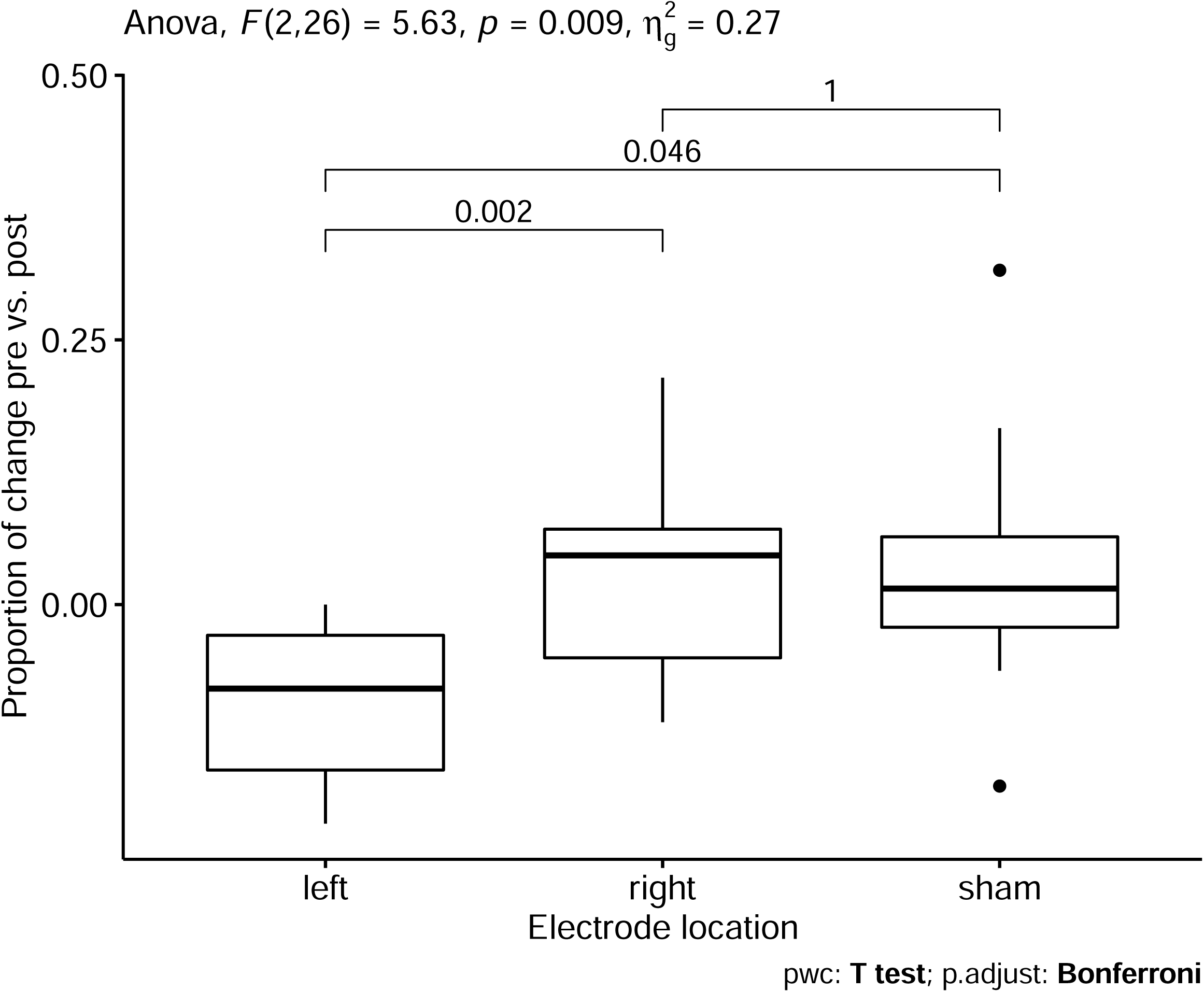
Repeated measures ANOVA results for percentage change after cathodal stimulation. After correcting for multiple comparisons, left hemisphere stimulation is still significantly lower than right hemisphere or sham stimulation.

We found that cathodal stimulation over the left SMG had a negative effect on pitch memory performance, while sham stimulation and cathodal stimulation over the right SMG did not affect performance. Furthermore, the proportion of change due to cathodal tDCS over the left SMG was significantly lower compared to either sham tDCS, or cathodal tDCS over the right SMG. In contrast, there was no difference between the effects of sham and cathodal tDCS over the right SMG. Performance at baseline did not differ between either of the active tDCS conditions and sham. These data provide evidence that applying cathodal tDCS over the left SMG led to a significant decrease in pitch memory performance, and that this effect was specific to the left hemisphere.

### 3.2 Experiment 2: Anodal tDCS - controlling for perceptual domain

We entered the pitch memory and visual memory data into separate 2 by 2 repeated-measures ANOVAs. The dependant variable was the total number of correct trials during task performance. The factors of the ANOVA were tDCS condition (sham, anodal), and time (pre-stimulation, post-stimulation).

#### 3.2.1 Pitch memory data

With regard to the pitch memory data, the ANOVA revealed a significant main effect of tDCS condition (*F(1,11) = 7*.*41, p = 0*.*02*). The active tDCS condition, with anodal stimulation over the left SMG, was associated with higher scores overall compared with sham according to a post-hoc comparison with Bonferroni correction. There was no main effect of time (*F(1,11) = 2*.*15, p = 0*.*17*). There was a significant interaction between the factors time and tDCS condition (*F(1,11) = 6*.*50, p = 0*.*03*), indicating that the difference between pre- and post-stimulation performance depended upon the tDCS condition. We ran a Bonferroni-corrected post-hoc test to determine if performance before stimulation differed from performance after stimulation for either of the tDCS conditions, and to compare baseline (pre) and post-stimulation performance across the two tDCS conditions (Fig. 5). The comparisons of pre- and post-stimulation performance for each stimulation condition revealed that only anodal stimulation over the left SMG had a significant effect (*F(1,11) = 18*.*4, p = 0*.*002*); there was a significant improvement in performance due to applying anodal tDCS over the left SMG. Pre- and post-stimulation performance did not differ for sham (*F(1,11) = 1*.*2, p = 0*.*59*). Baseline (pre) performance was the same for sham and active tDCS (*F(1,11) = 0*.*44, p > 0*.*9*). This baseline comparison established sham as a valid control condition for anodal tDCS over the left SMG. Post-stimulation performance did differ between anodal and sham stimulation (*F(1,11) = 13*.*7, p = 0*.*008*) as would be expected, with anodal stimulation corresponding to better performance.

**Fig 5:**
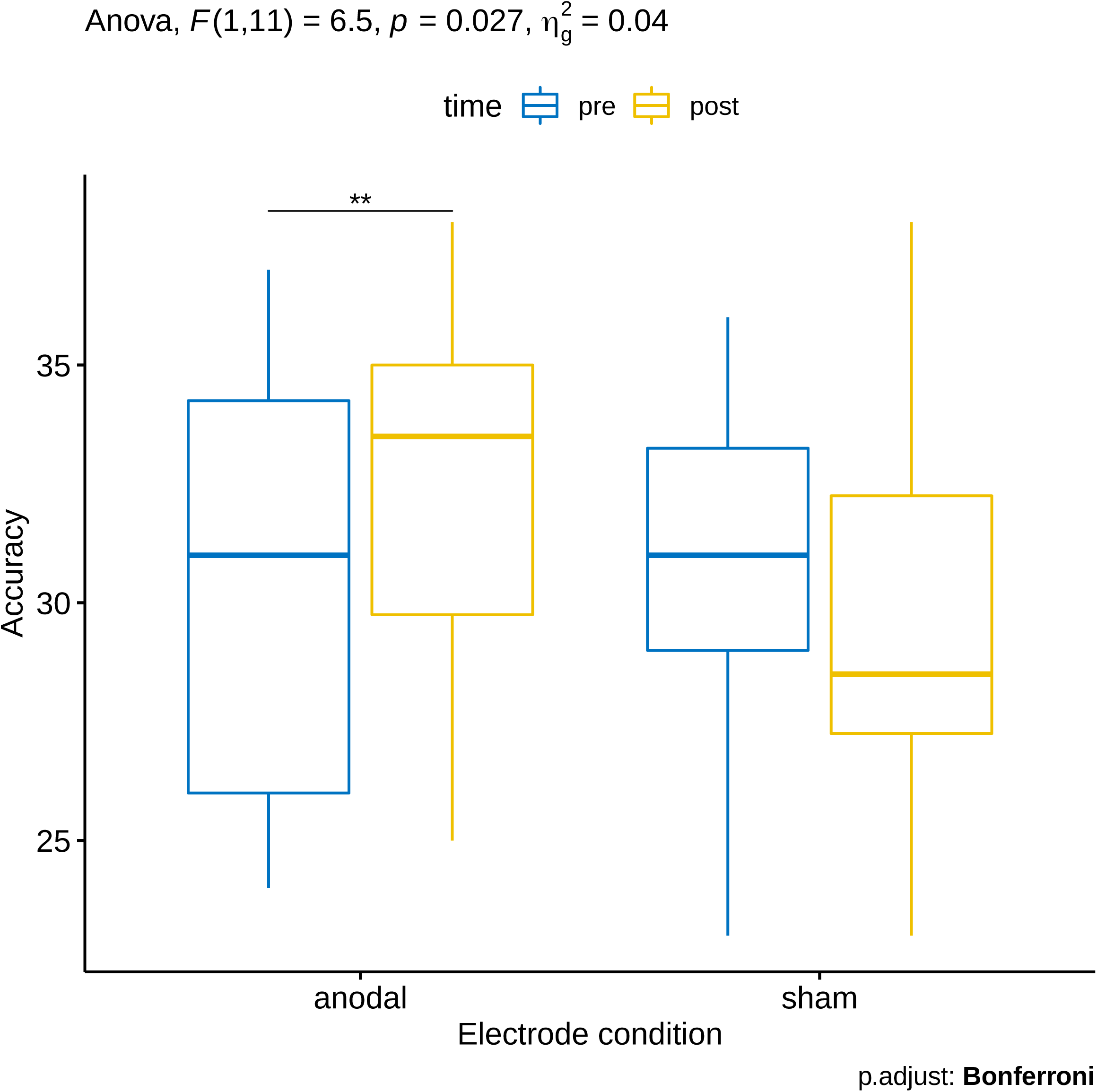
Repeated measures ANOVA and post-hoc comparisons revealed a main effect of electrode condition and an interaction between the factors electrode condition and time (pre vs. post) in the case of the pitch memory experiment, with a post-hoc test indicating that accuracy was significantly different for anodal stimulation but not sham.

We also calculated the proportion of change in performance from before to after tDCS (see Fig. 6), and entered the result into a paired-samples t-test comparing active tDCS and sham. This analysis yielded a significant result (*t(11) = 2*.*54, p = 0*.*03*), which established that there was a difference between the effects of sham and active tDCS. Stimulating the left SMG with anodal stimulation led to a significant improvement in performance compared to sham. Notably, the improvement in performance was mainly due to subjects performing below the mean at baseline, while those subjects performing above the mean showed either no change or negative change.

**Fig 6:**
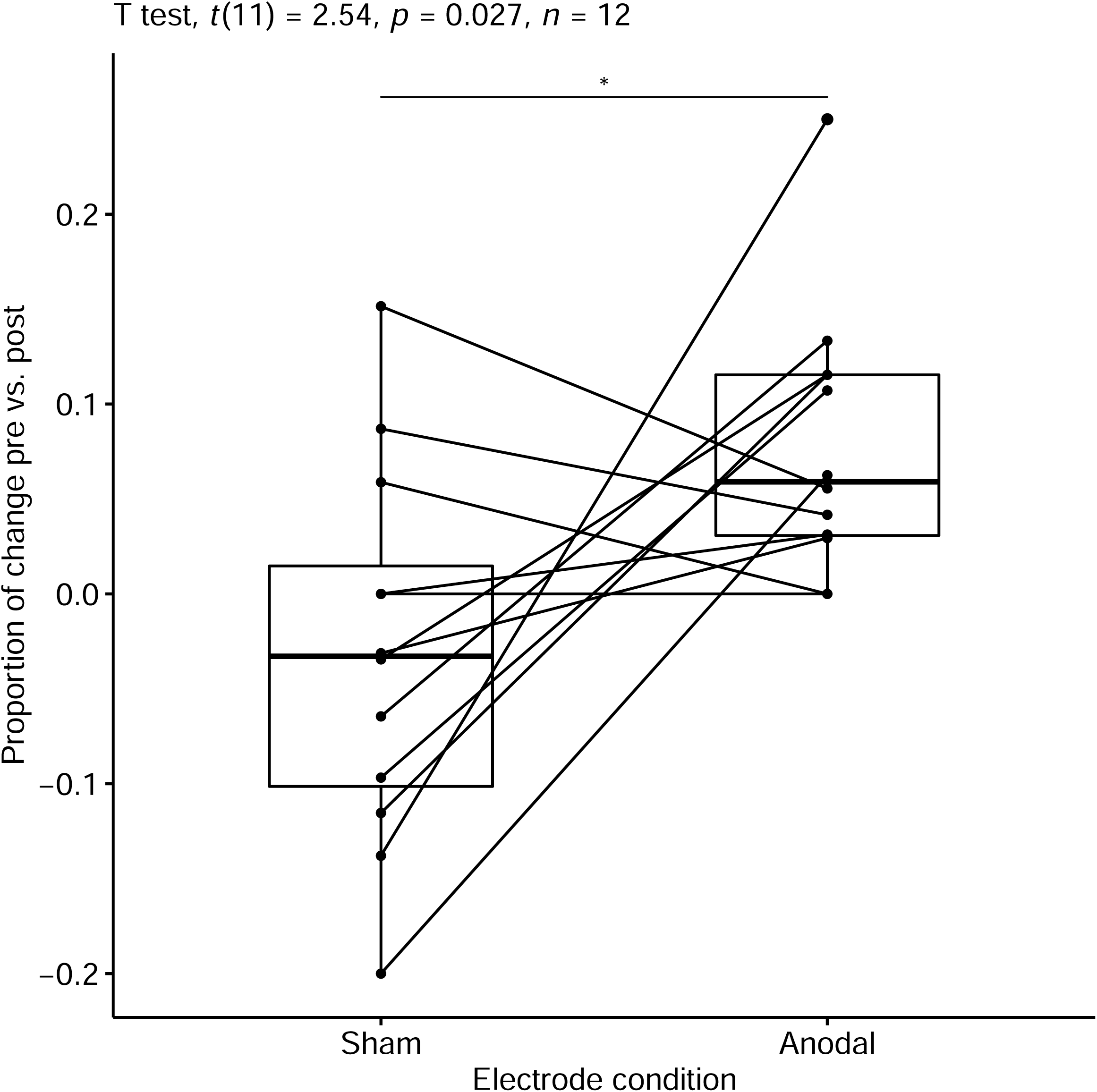
Individual change profiles and group results; paired-samples t-test showing significant difference in proportion of correct answers between sham and left anodal stimulation.

#### 3.2.2 Visual memory data

For the visual memory data, the 2 by 2 repeated-measures ANOVA yielded neither a significant main effect of time (*F(1,11) = 1*.*51, p = 0*.*24*), nor of tDCS condition (*F(1,11) = 2, p = 0*.*19*). There was also no significant interaction between the two factors, (*F(1,11) = 0*.*33, p = 0*.*58*). Baseline performance was the same for sham and active tDCS (*t(11) = 0*.*98, p = 0*.*35*). Therefore, the effect of real tDCS was no different from the effect of sham tDCS for the visual memory task.

The paired-samples t-test on the proportion of change in performance from before to after tDCS also revealed no difference between the two conditions (*t(11) = 0*.*69, p = 0*.*51*). This provided further evidence that the effects of active tDCS did not differ from sham for the visual memory task.

Notably, the baseline scores for the visual memory task (pre-sham: mean = 29.33, SD = 3.58, and pre-anodal: mean = 30.08, SD = 3.6) were similar to those for the pitch memory task (pre-sham: mean = 30.83, SD = 3.71, and pre-anodal: mean = 30.33; SD = 4.68). Because the visual memory task was of a similar design and of similar difficulty to the pitch memory task, it was a valid control for testing the domain specificity of the tDCS effects.

## 4 Discussion

We found that applying tDCS over the left SMG significantly affected pitch memory performance. The direction of the effects depended upon the polarity of the tDCS current: cathodal stimulation led to a deficit in performance, whereas anodal tDCS resulted in an improvement in performance. We used sham stimulation, which did not affect performance, to control for any placebo effect of the stimulation. We controlled for the location of stimulation by measuring the effects of applying tDCS over the right SMG, and did not find any effects on pitch memory with right SMG stimulation. This suggests that pitch memory processing is more lateralized to the left SMG. We also controlled for task specificity by measuring performance on a visual memory task, which tDCS did not affect. Only performance on the pitch memory task changed significantly. These data provide evidence that the left SMG is an important nodal point for short-term pitch memory, that the involvement of the SMG in pitch memory is lateralized to the left hemisphere, and that its contribution is unique to auditory short-term memory as opposed to visual memory.

This study provides further evidence for the causal role of the left SMG in short-term pitch memory, and supports the hypothesis that this region contributes more generally to short-term auditory working memory (1,2,6–9,25,30,47), including memory for both linguistic and non-linguistic sounds. It may be that the left SMG is a general auditory processing and short-term auditory storage module that is utilized for phonological and non-phonological processing, or that it is a language-devoted neural module that has general domain auditory processing potential (8). Further studies will need to compare short-term memory processing for pitch and phonological sounds, and the contribution of the left SMG to these operations. Interestingly, stimulation targeting Heschl’s gyrus or primary auditory cortex have shown that both right and left hemisphere play a role in pitch discrimination, but that the right Heschl’s gyrus seems to be more affected by cathodal stimulation (22,24). On the other hand rapidly changing acoustic cues seem to have more of a left hemisphere advantage (48) .

Only a limited number of studies have explored the potential for tDCS to have a negative or positive effect on cognitive performance, depending upon the polarity of stimulation; most of these studies have targeted prefrontal brain regions (27,49–51). Our results demonstrate the potential for tDCS to modulate cognitive performance in a task-specific, and polarity-dependent manner. Our results also highlight the value of multimodal approaches to studying brain-cognition relations. Non-invasive brain stimulation methods, such as tDCS and TMS, are useful tools for testing hypotheses derived from EEG, MEG, fMRI, or PET neuroimaging studies (52). Building upon our previous study (8), the present research shows how tDCS can be used to probe causal relations in a network of brain regions that are activated by a given cognitive task.

We hypothesize that the anodal tDCS that modulated excitability in the left SMG may have altered synaptic function by activating a mechanism related to long-term potentiation (53). This would account for the change in performance on the pitch memory task. The effects on performance depended upon the direction of the shift in excitability: increased excitability, due to anodal tDCS, improved performance, whereas decreased excitability, due to cathodal tDCS, had a negative impact. Shifts in cortical excitability in the human brain are the foundation of learning processes. Research has shown that the effects of tDCS on cortical excitability are dependent on voltage-gated pre- and post-synaptic sodium and calcium ion channels (54). Pharmacological studies have revealed that the efficacy of NMDA receptors mediates tDCS effects as well (55,56). (55) formed the following hypothesis regarding how anodal tDCS could lead to increased synaptic strength and learning: By increasing the resting firing rate of stimulated neurons, anodal tDCS may increase pre-synaptic activity, which facilitates membrane depolarization. The depolarization augments synaptic strength by means of NMDA receptor activity, most likely through an increase in intracellular Ca^2+^ levels. Cathodal tDCS would have the opposite effect. Therefore we hypothesize that the effect of tDCS on cognitive performance involving working memory is most likely mediated by NMDA receptor activity, for which there is evidence already in the literature (57–59). The findings of this study may be relevant to research on learning and learning disabilities, and to the study of tDCS modulation of cognitive functions in general.

## Acknowledgments

This work was supported by grants from the National Institute of Neurological Disorders and Stroke to BWV (NS053326) and from the National Institute of Deafness and Communications Disorders to GS (DC009823-01, DC008796, DC008796-02S1). BWV acknowledges support from the Grammy Foundation and from the Michael Smith Foundation for Health Research. We are grateful for Dr. Nora Raschle’s many contributions to this research study. Dr. Vines’s current affiliation is with Berklee Online, Berklee College of Music, Boston, MA. All authors declare no conflicts of interest.

